# The voltage-gated Ca_v_ Ca^2+^ channel subunit α_2_δ-4 is required for locomotor behavior and sensorimotor gating in mice

**DOI:** 10.1101/2022.01.17.476669

**Authors:** Annette Klomp, Ryotaro Omichi, Yoichiro Iwasa, Richard J. Smith, Yuriy M. Usachev, Andrew F. Russo, Nandakumar Narayanan, Amy Lee

**Author notes:** Department of Otolaryngology-Head and Neck Surgery, Okayama University Graduate School of Medicine, Dentistry and Pharmaceutical Sciences, Okayama 700-8558, Japan (Ryotaro Omichi). Department of Otorhinolaryngology, Shinshu University School of Medicine, Matsumoto 390-8621, Japan (Yoichiro Iwasa, Amy Lee). Dept. of Neuroscience, University of Texas-Austin, Austin, TX, 78712. Corresponding author: Amy Lee, Dept. of Neuroscience, University of Texas-Austin, 100 E. 24^th^ St., Austin, TX 78712.

## Abstract

Voltage-gated Ca_v_ Ca^2+^ channels are critical for the development and mature function of the nervous system. Variants in the *CACNA2D4 gene* encoding the α_2_δ-4 auxiliary subunit of these channels are associated with neuropsychiatric and neurodevelopmental disorders. α_2_δ-4 is prominently expressed in the retina and is crucial for vision, but extra-retinal functions of α_2_δ-4 have not been investigated. Here, we sought to fill this gap by analyzing the behavioral phenotypes of α_2_δ-4 knockout (KO) mice. α_2_δ-4 KO mice (both males and females) exhibited significant impairments in prepulse inhibition that were unlikely to result from the modestly elevated auditory brainstem response thresholds. Whereas α_2_δ-4 KO mice of both sexes were hyperactive in various assays, only females showed impaired motor learning/coordination in the rotarod assay. Female but not male α_2_δ-4 KO mice exhibited anxiolytic and anti-depressive behaviors in the elevated plus maze and tail suspension tests, respectively. Our results reveal an unexpected role for α_2_δ-4 in cognitive and motor function and identify α_2_δ-4 KO mice as a novel model for studying the pathophysiology associated with *CACNA2D4* variants.

## Introduction

Voltage-gated Ca_v_ Ca^2+^ channels mediate Ca^2+^ signals that initiate a vast array of signaling events including gene transcription, protein phosphorylation, and neurotransmitter release. The main properties of these channels are determined by a pore-forming α_1_ subunit, while auxiliary *β* and α_2_δ subunits regulate the trafficking and some functional aspects of these channels (1). These subunits are encoded by four genes each (2), with additional functional diversity conferred by extensive alternative splicing (3). The physiological importance of Ca_v_ channels is reflected in the numerous diseases that linked to mutations in the genes encoding the Ca_v_ subunits which include migraine, ataxia, and disorders of vision and hearing (4, 5).

In recent years, variants in Ca_v_ encoding genes have been consistently identified in genome-wide association studies of neuropsychiatric disorders. One of the most prominent of such studies analyzed single-nucleotide polymorphisms (SNPs) in ∼60,000 individuals and uncovered *CACNA1C*, the gene encoding Ca_v_1.2, as a major risk gene for schizophrenia, bipolar disorder, major depressive disorder, autism spectrum disorder, and attention deficit hyperactivity disorder (ADHD). Pathway analysis further revealed an association of other Ca_v_-encoding genes with these disorders, including *CACNA2D4* that encodes the α_2_δ-4 subunit (6). This result was rather unexpected given that α_2_δ-4 was thought to be expressed primarily in the retina, where it associates with the Ca_v_1.4 channel and regulates the structure and function of photoreceptor synapses (7-9).

α_2_δ is an extracellular protein that regulates the cell-surface trafficking of Ca_v_ channels (10), but may have additional roles. For example, α_2_δ-1 binding to thromobospondins promotes synapse formation in a manner that is inhibited by the analgesic and anti-convulsant drug, gabapentin (11). At the *Drosophila* neuromuscular junction, α_2_δ-3 is required for proper synapse morphogenesis—a process that does not involve its association with the Ca_v_2.1 channel (12). In the retina, the formation of photoreceptor synapses involves the role of α_2_δ-4 as a Ca_v_1.4 subunit and as a mediator of trans-synaptic interactions of the cell adhesion molecule, ELFN-1, with postsynaptic glutamate receptors (9).

Despite the association of α_2_δ-4 with neuropsychiatric disease, how α_2_δ-4 contributes to cognitive and affective functions is unknown. To address this question, we examined the behavioral phenotypes of α_2_δ-4 knockout (KO) mice (8).

## Materials and Methods

### Animals

All procedures using animals were approved by the University of Iowa Institutional Animal Care and Use Committee. The α_2_δ-4 KO mouse line was bred on a C57/Bl6 background and characterized previously (8). Separate cohorts of males (15-25 week old, n= 10 wild-type (WT), n= 11 KO) and females (11-22 week old, n= 11 WT, n= 11 KO) were analyzed. Before beginning handling and testing, mice were ear punched for identification. All mice were housed in groups of 2-3 animals per cage for the duration of the handling and testing periods with food and water ad libitum (describe details of the cages). The room in which the mice were housed was maintained on a consistent light cycle with lights on at 0900 and lights off at 2100 and testing took place between 0800 to 1300. Males and females were tested in separate cohorts at different time points to prevent pheromones on the testing apparatus from impacting results. Mice were generally acclimatized for 30 min in the room in which the assay was conducted prior to initiating the test. A full week was taken between every test to reduce the impact of stress from previous tests on the next result. The order of testing was designed to minimize the impact of preceding assays by performing those with the least stressful tasks first.

### Auditory Brainstem Responses

Auditory brainstem responses (ABRs) were performed as described previously (13). Mice were anesthetized with intraperitoneal injection of ketamine (100 mg/kg) and xylazine (10 mg/kg). Recordings were conducted on both ears of all animals on a heating pad using electrodes placed subcutaneously in the vertex and underneath the left or right ear. Clicks were square pulses 100 ms in duration, and tone bursts were 3 ms in length at distinct 8-, 16-, and 32 kHz frequencies. ABRs were measured using BioSigRZ software (Tucker-Davis Technologies), with stimulus levels adjusted in 5-dB increments between 25 and 100 dB SPLs in both ears. Electrical signals were averaged over 512 repetitions and ABR threshold was defined as the lowest sound level at which a reproducible waveform was measured.

### Elevated Plus Maze

The testing apparatus consisted of a plus-shaped maze elevated 40 cm above the floor. Two opposing closed and open arms extended from a central zone. Open arms had no walls whereas closed arms were surrounded by gray walls. The floor of the maze was made of gray plastic material. Illumination intensity in the central square was approximately 500 lux. Mice were moved from the home cage to the central square of the maze, always facing the same closed arm. The animals were allowed to explore the maze for 10 min. In the event of a fall, the animal was placed in the central square facing the same closed arm and recording resumed. Time spent in the open and closed arms was evaluated using video recording and Anymaze software.

### Light Dark Box

The testing apparatus consisted of a chamber divided into a light and dark compartment equipped with infrared beam tracking (Med Associates). The apparatus was divided into 2 chambers with a gap in the wall between them. Mice were tested using a very bright light in the light chamber (27,000 lux). Mice were moved from the home cage to the light side of the apparatus facing away from the dark chamber. The animals were allowed to freely explore and move between the chambers for 30 min and the animals’ movements were documented in sequential 5 min intervals via infra-red tracking. Time spent in either compartment was analyzed by Activity Monitor software.

### Open Field Test

The testing apparatus consisted of an open square chamber with walls of 40 cm height and width. Illumination intensity in the central square was approximately 500 lux. Mice were moved from the home cage to the center of the open chamber. The animals were allowed to freely explore the chamber for 10 minutes. Animal behavior was evaluated using video recording and Anymaze software. Relative time spent in the inner and outer portion of the box were taken as a measure of the animals’ anxiety-like behavior. Total distance traveled over the 10 minutes was taken as a measure of the animals’ basal activity level.

### Prepulse Inhibition

The testing apparatus consisted of a startle response box (SR-LAB from San Diego Instruments). A restraint chamber consisted of a clear plastic tube from which the tremble response of the animal could be measured via an accelerometer underneath the chamber. Animals were placed in the restraint chamber and allowed to acclimate to the chamber for 10 min with a consistent background white noise level of 65 dB which was present for the entire experiment. The 25-min testing period was divided into 3 blocks each consisting of 6 or 60 trials. All trials were presented with a randomly spaced intertrial interval ranging from 7 to 15 seconds. The first block consisted of 6 pulse trials at 120 dB. The second block contained 12 of each of the following trial types: standard pulse at 120 dB, no stimulation, prepulse of +4 dB above background, prepulse of +8 dB, and prepulse of +16 dB. The third block consisted of 6 pulse trials at 120 dB. Startle response amplitudes (in mV) were measured in SR-LAB software and %PPI measured as (startle response for pulse alone - startle response for pulse with pre-pulse) / startle response for pulse alone) X 100.

### Rotarod Test

The testing apparatus consisted of a rotating spindle 3.0 cm in diameter that will increase in speed over the course of the trial (Rotamex 5). Mice were trained for 2 consecutive days with 3 testing trials per mouse each day separated by at least 30 min. For the testing trial, the speed of rotation was increased by 1.2 rpm every 20 s and the latency to fall was recorded. The 6 testing trials were averaged for each mouse.

### Forced Swim Test

The testing apparatus consisted of a 2-liter beaker filled with 1200 ml of water at room temperature. Mice were placed in the water and monitored for 6 min, then were dried and placed in a recovery cage with a cage warmer. Time spent immobile was recorded, with immobility defined as lack of motion in the hind legs except necessary movement to balance and keep the head above the water.

### Tail Suspension Test

The testing apparatus consisted of a metal bar suspended 30-40 cm above the table. Tails of the mice were wrapped in adhesive tape within the last 1 cm of the tail. A clear plastic tube was placed around the animal’s tail to prevent climbing up the tail and onto the bar. Time spent immobile was recorded, with immobility defined as lack of attempting to move their limbs as described previously (14).

### Statistics

Statistical analysis was done with GraphPad Prism software 8.0 and RStudio. An alpha level of 0.05 was used for all statistical tests. For datasets without repeated measures, data were first tested for normality by the Shapiro–Wilk test and homogeneity by Levene’s test. For parametric data, ANOVA with post hoc Holm-Sidak’s multiple comparisons test was performed. For non-parametric data, Kruskal Wallis tests were used with post hoc Dunn’s multiple comparisons. For data sets with repeated measures, a repeated measures linear mixed model was used with post hoc estimated marginal means. The main effects were reported if there was no significant interaction, and post hoc analysis was performed on the main effects that had more than two levels. Otherwise, post hoc tests were performed and simple main effects were reported using adjusted *p* value for multiple comparisons. Data were graphically represented as mean ±standard error of the mean (SEM) for each group. Results were considered significant when *p* < 0.05 (denoted in all graphs as follows: \**p* < 0.05; \*\**p* < 0.01; \*\*\**p* < 0.001).

## Results and Discussion

α_2_δ-4 KO mice were born at normal Mendelian ratios and did not exhibit any overt behavioral phenotypes other than hyperactivity. The control wild-type (WT) strain corresponded to C57Bl6 strain on which the α_2_δ-4 KO mice were bred for at least 10 generations. Cohorts of male and female mice were analyzed separately, and there were no differences in body weight of the WT and α_2_δ-4 KO mice used in this study (Table 1).

**Table 1:** Body weights (g) of animal subjects in this study

| Cohort: | WT F | WT M | KO F | KO M |
| --- | --- | --- | --- | --- |
|  | 22 | 26.1 | 24.4 | 31.1 |
|  | 23.4 | 25.1 | 22.1 | 28.7 |
|  | 20.7 | 27.5 | 21.5 | 29.4 |
|  | 19.9 | 35.1 | 27.8 | 33 |
|  | 23.4 | 30.4 | 26.6 | 29.9 |
|  | 21.3 | 32.9 | 22 | 27.6 |
|  | 22.8 | 33.1 | 20.7 | 31.6 |
|  | 24 | 28.4 | 20.4 | 27 |
|  | 22 | 29.2 | 20.7 | 30.9 |
|  | 18.5 | 27.7 | 21.7 | 25.8 |
|  | 20 |  | 19.8 | 27.1 |
| **Mean** | **21.636** | **29.55** | **22.518** | **29.282** |
| **StDev** | **1.6499** | **3.1004** | **2.4998** | **2.1485** |

Animals (n=43) were weighed prior to any of the behavioral tests in this study. There was no significant effect of genotype on body weight in cohorts of females (**F**; $F_{9,10} = 2.103$ , $p = 0.6857$ ) or males (**M**; $F_{1,39} = 7.876$ , $p = 0.8277$ ).

### Prepulse inhibition is impaired in α_2_δ-4 KO mice

Sensorimotor gating is a form of pre-attentive processing that is commonly studied in humans and animals using prepulse inhibition (PPI). In this test, a response to a strong acoustic stimulus is generally diminished when it is preceded by a subthreshold stimulus (15). Reductions in PPI are thought to reflect impairments in working memory in individuals diagnosed with schizophrenia, bipolar disorder, and post-traumatic stress disorder and in animal models of these conditions (16, 17). Because of the association of Ca_v_-encoding genes with these disorders (6), we tested whether α_2_δ-4 KO mice exhibit deficits in PPI. WT and α_2_δ-4 KO were tested for startle responses to a 120 dB acoustic stimulus that was administered alone or after a prepulse stimulus of 4, 8, or 16 dB, and PPI was expressed as the % change in the response amplitude due to the prepulse (%PPI, Fig.1A,B). In this assay, there was a significant main effect of both sex (*F*_*1, 39*_= 7.876, *p* < 0.01) and genotype (*F*_*1, 39*_= 10.26, *p* < 0.01), but no interaction between these variables (*F*_*1,39*_= 0.0028, *p* = 0.958; Fig.1B). PPI was significantly lower for α_2_δ-4 KO than for WT mice in the cohort of females (*p* < 0.05 for both 8 and 16 dB prepulse) and males (*p* < 0.05 for 8 dB, *p* < 0.01 for 16 dB prepulse). In some mouse strains, relatively low levels of PPI correlate with low basal startle amplitudes (17). However, basal startle amplitudes were significantly higher in α_2_δ-4 KO mice than in WT mice (*F*_*1,39*_= 55.50, *p* < 0.001; Fig.1C). Some studies have shown that patients with schizophrenia have an impaired habituation to the startle pulse (18), which would manifest as a difference in startle response to the 120 dB-stimulus administered without the prepulse (blocks 1-3, Fig.1A). There was no effect of genotype on this parameter (*F*_*2, 78*_= 1.580, *p* = 0.213). Collectively, these results show that α_2_δ-4 KO mice exhibit impaired PPI without alterations in habituation.

**Figure 1.**
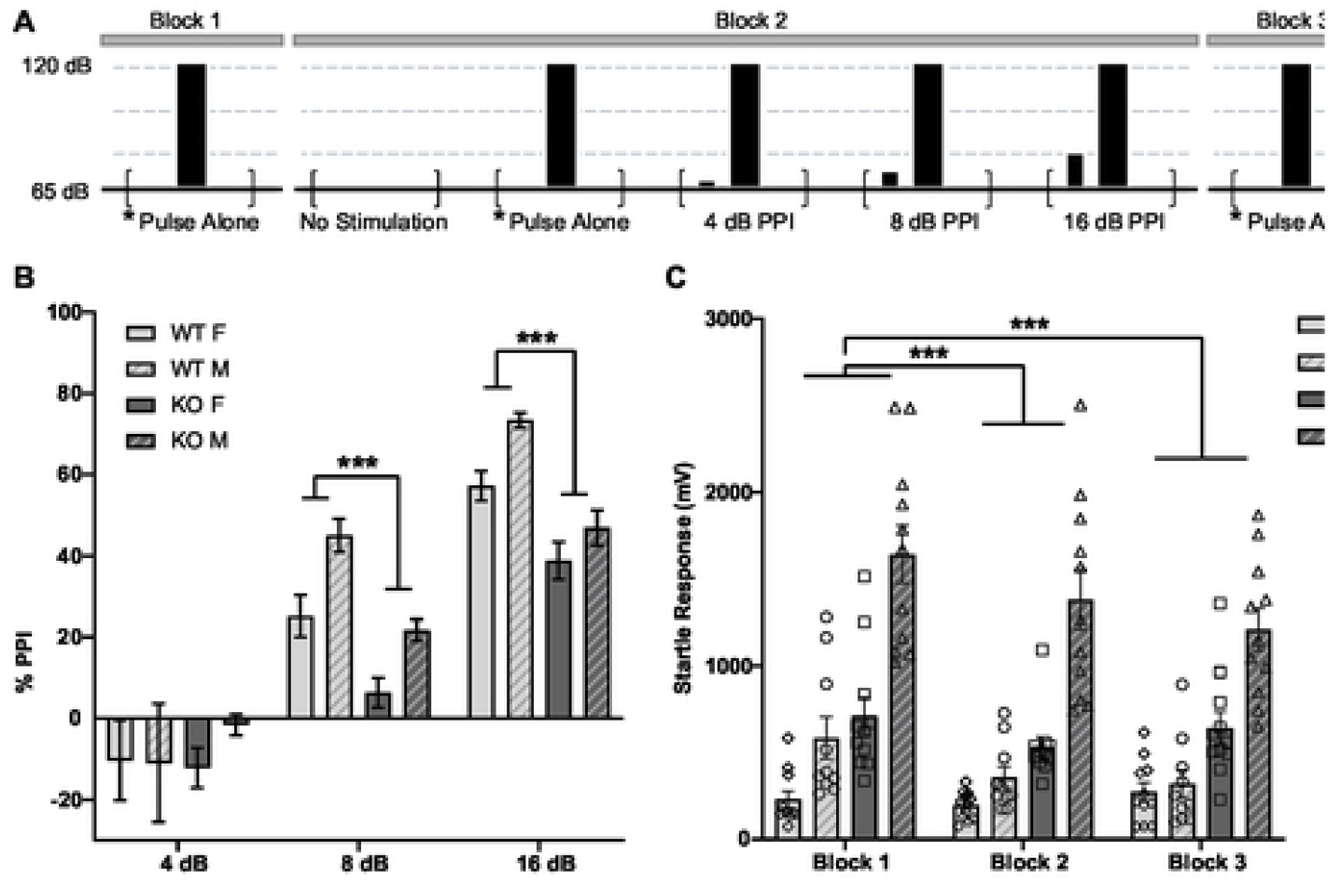
α_2_δ-4 KO mice exhibit impaired PPI. (**A**), Schematic of PPI session design. Block 1 & 3 each consist of 6 test pulses (120 dB) alone trials. In block 2, animals were exposed to a period of no stimulation, the test pulse alone, or test pulses preceded by a PPI prepulse of 4, 8, or 12 dB. Each trial type (total of 12) was presented in a randomized order with varying intertrial interval. Each test pulse was 40 ms and prepulse was 20 ms in duration. (**B**) %PPI evoked by the indicated prepulse intensities. (**C**) Startle response amplitudes evoked by test pulses without prepulses. \**p* < 0.05; \*\**p* < 0.01; \*\*\**p* < 0.001 by linear mixed model.

In the retina and cochlea, α_2_δ proteins support the activity of Ca_v_1.4 and Ca_v_1.3 channels that mediate glutamate release at the specialized ribbon synapse of photoreceptors and inner hair cells, respectively (19). Although detailed analyses of α_2_δ variants in the cochlea are lacking, α_2_δ-4 has been detected in cochlear hair cells by single cell RNA sequencing (20). To determine whether hearing impairment could contribute to weakened PPI in α_2_δ-4 KO mice, we measured auditory brain stem responses (ABRs). In this assay, elevated ABR thresholds correlate with hearing deficits. In response to click stimuli, α_2_δ-4 KO males had significantly higher thresholds than WT males (*p* < 0.05). For pure tone stimuli from 8 kHz to 32 kHz, there was no overall effect of genotype or sex (*F*_*1, 60*_= 1.0140, *p* = 0.318 & *F*_*1, 60*_= 2.3092, *p* = 0.134), but an interaction between genotype and sex (*F*_*1, 60*_= 8.0533, *p* < 0.01) indicated lower thresholds in α_2_δ-4 KO females than in WT females (*p* < 0.01, Fig.2). Nevertheless, all α_2_δ-4 KO mice displayed functional hearing above 60 dB, the range used in the PPI assays, which argues against the possibility that the reduced PPI of the α_2_δ-4 KO mice resulted from hearing loss.

**Figure 2.**
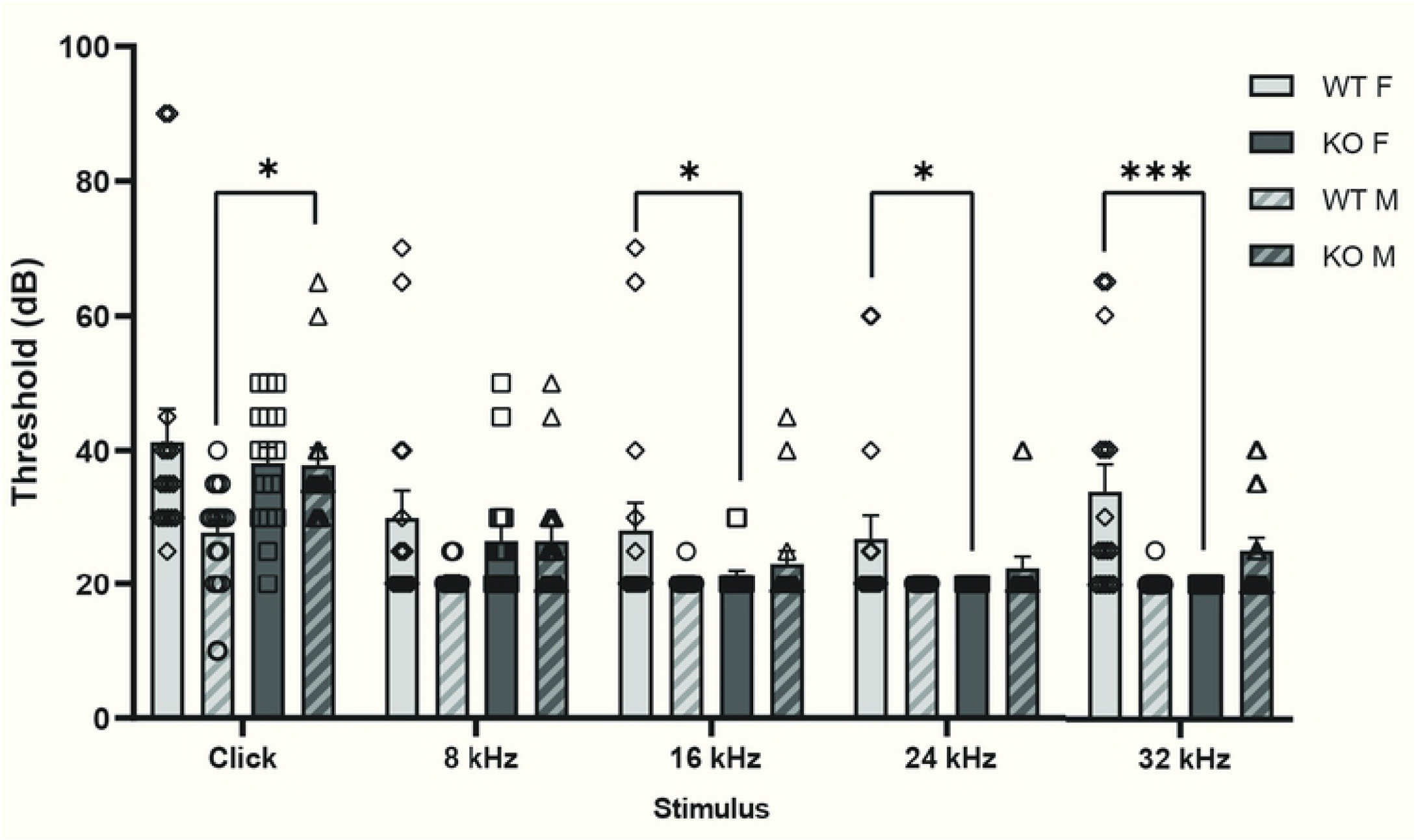
α_2_δ-4 KO mice exhibit sex-specific alterations in ABRs. Thresholds (dB) were plotted for WT and α_2_δ-4 KO mice in response to click and pure tone stimuli of the indicated frequencies. \**p* < 0.05; \*\**p* < 0.01; \*\*\**p* < 0.001 by Kruskal-Wallis, Dunn’s test, linear mixed model, and estimated marginal means.

### α_2_δ-4 KO mice exhibit anxiolytic and antidepressant phenotypes

Anxiety and depression are common features of a variety of neuropsychiatric disorders, including those associated with *CACNA1C* variants (21) and have a high rate of comorbidity with schizophrenia (22, 23), ADHD (24), ASD (25), bipolar disorder (23, 26), and major depressive disorder (23, 27). Therefore, we tested the performance of α_2_δ-4 KO mice in behavioral assays designed to assess anxiety (open field test, OFT; elevated plus maze, EPM; and light dark box, LD) and depression (forced swim test, FST; and tail suspension test, TST). In the OFT, the animals are placed in the center of an open chamber and the time spent avoiding the center is used as a metric for anxiety-like behavior (*i*.*e*., thigmotaxis). In the EPM, the animals are placed in the center of a raised platform with open and closed arms and the time spent avoiding the open arms is taken as an indicator of anxiety-like behavior. While α_2_δ-4 KO and WT mice did not differ in thigmotaxis in the OFT (*p* = 0.99, Fig.3A,B), α_2_δ-4 KO mice spent more time in the open arms of the EPM than WT mice (Open *η*^2^ = 0.141, *p* < 0.01 by Kruskal-Wallis; Closed *F*_*1, 34*_= 14.206, *p* < 0.001 by linear mixed model; Fig.3C,D). It is unlikely that visual impairment of the α_2_δ-4 KO mice influenced their abilities to respond to the aversive stimuli of the OFT and EPM since these mice have normal vision in daylight but not dim light conditions (8). As a further test, we performed the light dark box assay in which avoidance of a chamber with a bright light stimulus is taken as a measure of anxiety-like behavior. The α_2_δ-4 KO spent more time in the lighted chamber than WT mice (*η*^2^ = 0.100, *p* < 0.05 by Kruskal-Wallis; Fig.3E,F). The light intensity used in the lighted chamber was 27,000 lux, which is well above the visual threshold for α_2_δ-4 KO mice (8). Taken together, results from the EPM and LD assays support an anxiolytic phenotype in α_2_δ-4 KO mice.

**Figure 3.**
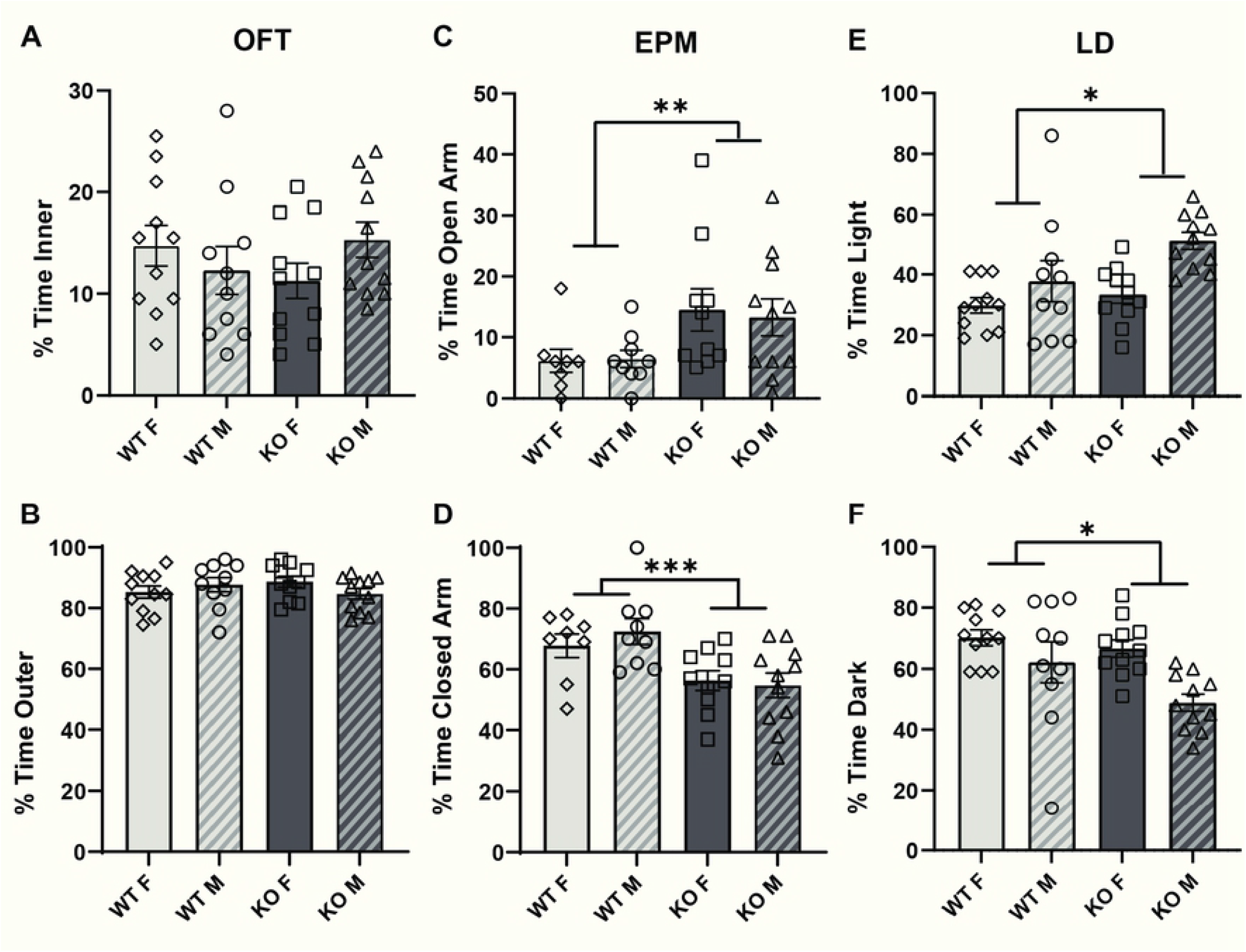
α_2_δ-4 KO mice exhibit diminished anxiety-like behaviors. For WT and α_2_δ-4 KO mice, graphs show the % total time spent in the inner and outer regions of the chamber in the open field test (OFT) (**A**,**B**), open and closed arms of the elevated plus maze (EPM) (**C**,**D**), and light and dark chambers in the light-dark assay (LD) (**E**,**F**). \**p* < 0.05; \*\**p* < 0.01; \*\*\**p* < 0.001 by Kruskal-Wallis and linear mixed model.

In the TST and FST, depressive phenotypes are measured as the duration of immobility following suspension of the animal by its tail, or placement of the animal in a beaker of water, respectively. While there were no differences between genotypes in the FST, α_2_δ-4 KO mice spent significantly less time immobile than WT mice in the TST (*F*_*3, 252*_= 15.04, *p* < 0.001; Fig.4A-F). These results indicate a task-specific antidepressant-like phenotype in the α_2_δ-4 KO mice.

**Figure 4.**
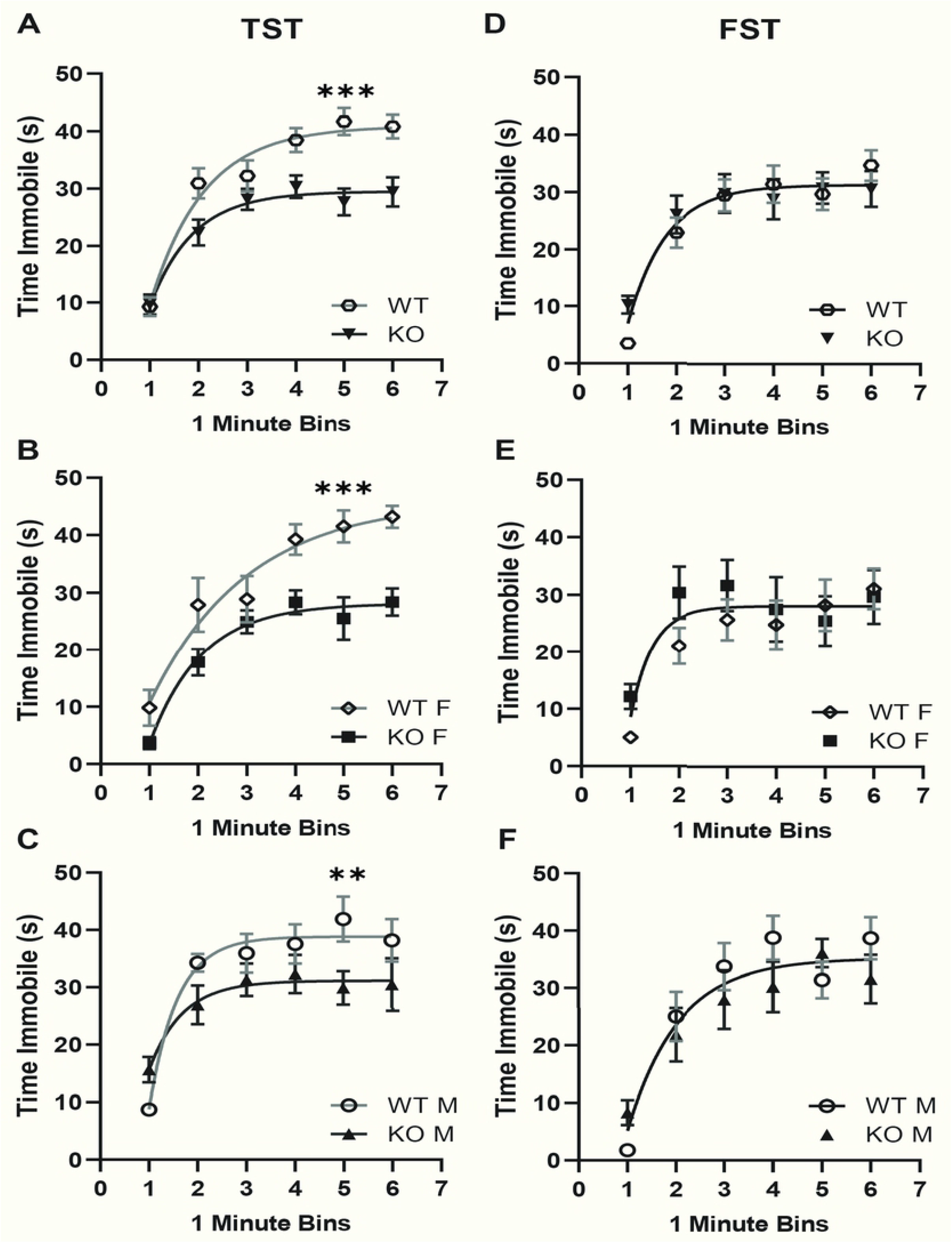
α_2_δ-4 KO mice exhibit diminished depression-like behavior in the tail suspension test. For WT and α_2_δ-4 KO mice, the duration spent immobile in the tail suspension test (TST) (**A-C**) and forced swim test (FST, (**D-F**) was plotted against time (in 1-min bins) during the assay. **A** and **D** represent results for males and females combined while **B**,**C**,**E**,**F** show data disaggregated by sex. Smooth line represents exponential fits of the results. \**p* < 0.05; \*\**p* < 0.01; \*\*\**p* < 0.001 by nonlinear regression.

### α_2_δ-4 KO mice exhibit abnormal motor behavior

Abnormal motor behaviors are a common feature of neurodevelopmental disorders including ASD and ADHD (28-30). Thereofre, we tested motor function of α_2_δ-4 KO mice in the rotarod assay. In this assay, the mice are placed on a rotating cylinder that is gradually accelerated and the length of time the animal can stay on the cylinder is taken as a measure of balance, coordination, and motor planning (31). The latency to fall was shorter for α_2_δ-4 KO than for WT mice (*F*_*1, 39*_= 6.457, *p* < 0.05; Fig.5A). To further assess motor phenotypes in the α_2_δ-4 KO mice, we analyzed data in the OFT, EPM, and LD assays for aberrant locomotion. In each case, the total distance traveled by α_2_δ-4 KO (both males and females) mice was significantly greater than for WT mice (OFT *η*^2^ = 0.266, *p* < 0.001; EPM: *F*_*1, 34*_= 16.09, *p* < 0.001; LD: *η*^2^ = 0.490, *p* < 0.001; Fig.5B-D). These results show that α_2_δ-4 KO mice exhibit signs of hyperactivity and sex-dependent impairment in motor coordination.

**Figure 5.**
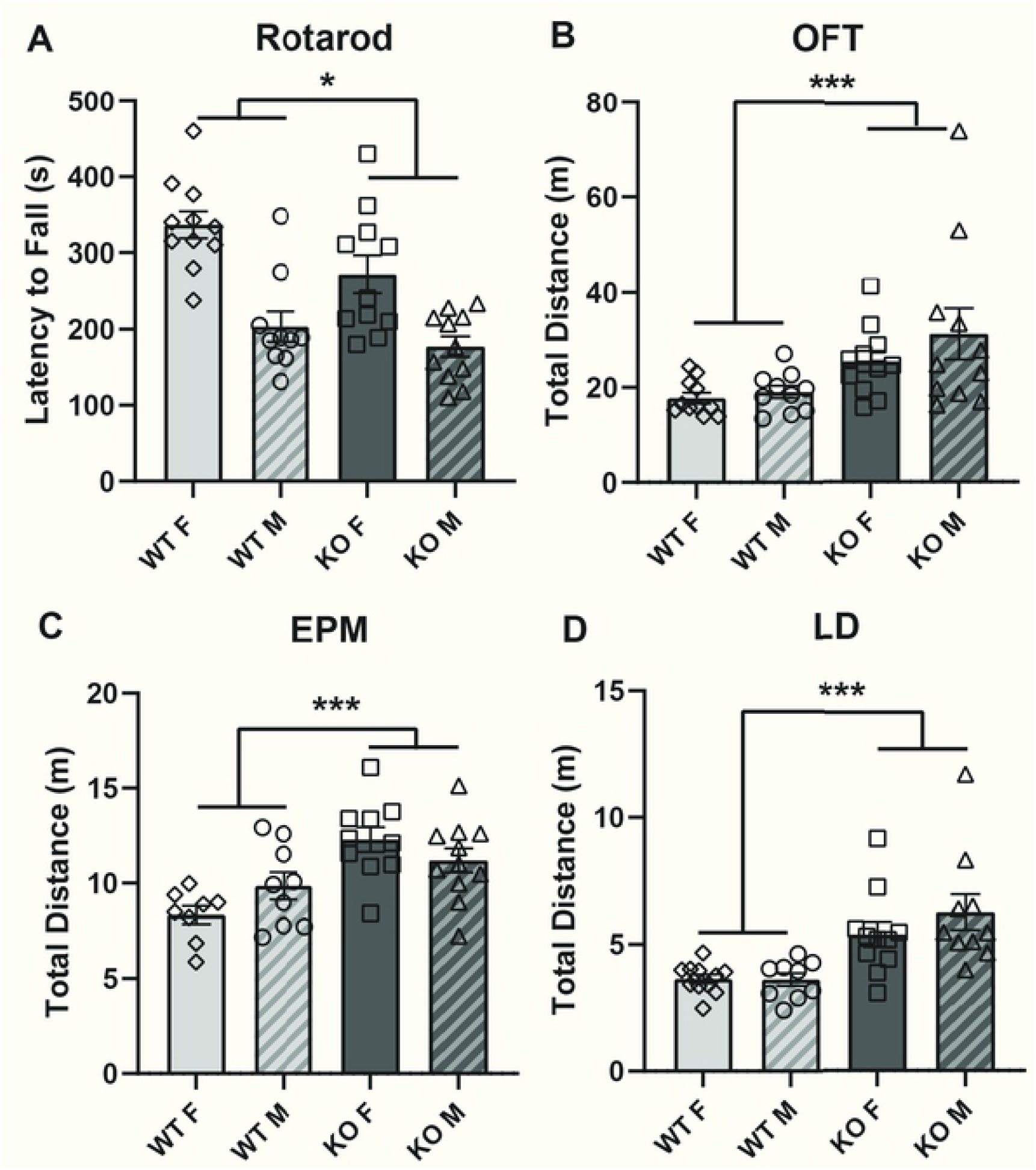
α_2_δ-4 KO mice exhibit alterations in motor behavior. For WT and α_2_δ-4 KO mice, graphs show the latency to fall in the rotarod assay **(A**) and total distance traveled in the OFT (**B**), EPM (**C**), and LD (**D**) assays. Rotarod: Genotype *F*_*1, 39*_= 6.457, *p* < 0.05; Sex *F*_*1, 39*_= 22.543, *p* < 0.001; Genotype:Sex *F*_*1, 39*_= 1.806, *p* = 0.1867 by one-way ANOVA. \**p* < 0.05; \*\**p* < 0.01; \*\*\**p* < 0.001 by Kruskal-Wallis and one-way ANOVA.

Our results show that α_2_δ-4 KO mice exhibit a pattern of cognitive, affective, and motor behaviors that resemble those in neuropsychiatric disorders that are linked to variants in Ca_v_-encoding genes (6). Because α_2_δ proteins enhance the cell-surface trafficking of Ca_v_ channels (5), the phenotypes of α_2_δ-4 KO mice could result from loss-of function of Ca_v_ channels in key brain regions such as the hippocampus, cerebral cortex, and cerebellum. A caveat is that compared to other α_2_δ subunits, α_2_δ-4 is nearly undetectable in these brain regions (32). Moreover, mice lacking the expression of the major Ca_v_ subtype implicated in neuropsychiatric/neurodevelopmental disorders (*i*.*e*., Ca_v_1.2) exhibit increased anxiety-like behavior α_2_δ-4 KO mice (33) whereas α_2_δ-4 KO mice present with an anxiolytic phenotype in the EPM and LD assays (Fig.3C-F). However, deletion of Ca_v_1.2 in the prefrontal cortex causes anti-depressant behavior in the TST (34), similar to that of α_2_δ-4 KO mice in our study (Fig.4A-C). α_2_δ-4 could be expressed in a small subset of neurons implicated in these behaviors, thus escaping detection by quantitative PCR in homogenates of specific brain regions (32). Alternatively, α_2_δ-4 could undergo pathological upregulation. For example, α_2_δ-4 expression is increased in hippocampus of humans and mice following epileptic seizures (35). Given that α_2_δ proteins regulate synapse formation in part through trans-synaptic interactions with proteins other than Ca_v_ channels (9, 11, 12, 36), aberrant expression of α_2_δ-4 could cause defects in neuronal connectivity that modify cognitive/affective behaviors.

Although blind under dim-light conditions, α_2_δ-4 KO mice are expected to have normal vision under the lighting conditions used in our study (8, 9). To date, alterations in cognitive and/or affective function in individuals diagnosed with *CACNA2D4-*related vision impairment have not been reported. However, the etiology of most neuropsychiatric disorders is complex and likely involves hundreds to thousands of risk alleles distributed across the genome (37). Our findings that α_2_δ-4 KO mice exhibit defects in PPI, motor function, and anxiety/depression-related behaviors validate the importance of *CACNA2D4* as one such risk allele and that studies of the extra-retinal functions of α_2_δ-4 warrant further study.

## Acknowledgements

The authors thank Jussara Hagen for expert technical assistance, Anjali Rajadhyaksha and Charlotte Bavley for helpful input on analysis of behavioral studies.

